# A fatigue-resistant myoneural actuator for implantable biohybrid systems

**DOI:** 10.1101/2025.03.14.642606

**Authors:** Hyungeun Song, Guillermo Herrera-Arcos, Gabriel N. Friedman, Seong Ho Yeon, Cassandra He, Samantha Gutierrez-Arango, Sapna Sinha, Hugh M. Herr

## Abstract

Implantable biohybrid systems with computer-controlled actuation offer the capacity to modulate biological forces, but require biocompatible, self-sustaining, and scalable actuators. Repurposing biological muscles can fulfill this need. However, muscle fatigue limits the fundamental capabilities of muscle-actuated systems. Here we present a fatigue-resistant myoneural actuator (MNA) with engineered recruitment biophysics in a rodent model. The MNA is based on manipulating native axonal composition through sensory reinnervation. This regenerative approach redirects volitional control to computer control via nerve stimulation while maintaining self-sustainability. Compared to native muscles without the myoneural manipulation, fatigue resistance is augmented by 260%. Furthermore, we demonstrate closed-loop control with reversible neural isolation of the actuator, preventing unintended neural signaling to the central nervous system during operation. To illustrate the potential of the MNA technology, we present a biohybrid neuroprosthetic interface and a biohybrid organ system capable of modulating neural afferents and organ mechanics, respectively. Our framework demonstrates augmented biological muscle actuation while maintaining inherent tissue properties, bridging the technological gap for implantable biohybrid systems.

## Main

Actuation is integral to living systems, governing cellular, tissue, and organ functionality^1–3^. Being able to actuate living systems in a controllable manner can regulate biological mechanisms underlying their function^4–9^. Implantable biohybrid systems with computer-controlled actuation is a promising approach to modulate human biology. For example, actuation of muscle-tendon stretch could modulate neural afferents for limb proprioception, enabling neural feedback from a prosthetic limb or a virtual avatar (Fig. 1a). Similarly, controlling intestinal actuation could restore failed organ function or modulate the gut-brain axis, enabling physiological feedback such as appetite through neural afferents (Fig. 1a). Biohybrid systems require biocompatible, self-sustaining and controllable actuators, while also requiring design flexibility to interface with various organ systems across scale. Although several critical advances have been made in synthetic actuator and biofabrication technologies, engineering science has not yet produced an actuator that can fully meet these demands^10–13^.

**Fig. 1.**
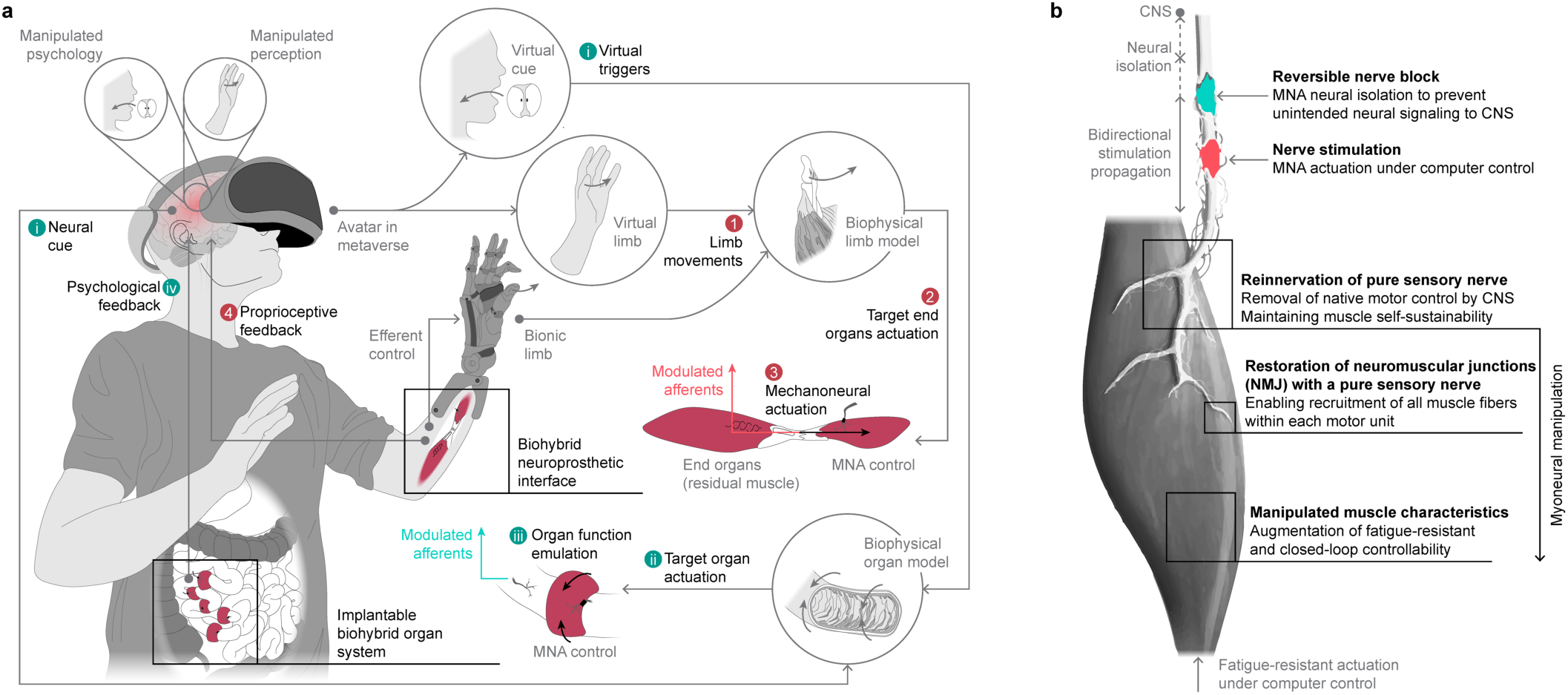
Myoneural actuator (MNA) for biohybrid systems. **a**, The ability to control organ actuation facilitates modulation of human biology. For example, actuation of muscle-tendon state and force through a serially-coupled MNA allows neural afferent modulation responsible for limb proprioception, enabling afferent feedback for bionic or virtual limbs (loop 1-4). Furthermore, MNAs could emulate organ mechanics, such as intestinal contraction, driven by neural or virtual cues, thereby assisting failed organ function and offering physiological feedback, such as appetite modulation (loop i-iv). **b**, Components of the fatigue resistant MNA. To transform a native muscle into a MNA, we denervate its motor nerve and reinnervate it with a pure sensory nerve, thereby redirecting motor control from the central nervous system (CNS) to a computer-controlled framework. Electrical nerve stimulation modulates the MNA, with reversible nerve block applied proximally to neurally isolate the MNA from the CNS during operation. The proposed myoneural manipulation serves as an enhanced interface for motor recruitment, rendering it a fatigue-resistant actuator with improved closed-loop control performance.

Linear synthetic actuator technologies such as electromagnetic, pneumatic or hydraulic actuators have the limitations of being heavy and large, and together with their battery-powered activation systems, cannot be easily reduced to the centimeter size scale^11,13^. Additionally, their material composition makes them inherently non-biocompatible^13^. Conversely, tissue-engineered muscle actuators leverage the unique properties of muscle cells, including nanometer-scale molecular machinery regulated by ion control for contraction^11^. However, such actuators face challenges such as complex tissue engineering techniques for 3D cellular organization, maintaining a continuous biochemical supply, integration of vascular and neural networks, and limited scalability^14,15^.

As a potential resolution to these difficulties, repurposing skeletal muscle as a viable actuator for biohybrid applications offers several critical advantages. Muscle tissue possesses inherent biocompatibility and self-sustainability^11^. Muscle has impressive energetics due to metabolic efficiencies, a scalability from micrometer to meter scales, and a neural innervation that enables high-fidelity control^10,16^. Muscle exhibits functional adaptation to environmental demands through hypertrophic and hyperplastic growth and is capable of self-repair in case of damage^10^. Moreover, muscle provides an adaptive compliant media with a form factor that can be tensioned, wrapped, or rolled to interface with various organs^17,18^. Incorporating a minimal synthetic component, such as a muscle or nerve electrode, enables computer-controlled actuation under functional electrical stimulation (FES). However, a critical limitation of FES is muscle fatiguability during continuous electrical activation, limiting the fundamental capability of muscle-actuated systems^19,20^.

Various external interventions to extend muscle fatigue resistance beyond its natural form have been explored, including chemical dosing, advanced electrode design, dynamic stimulation protocols, and novel stimulation modalities^20–23^. These approaches mainly focus on constructing rich environments for glucose metabolism, engineering synthetics, and genetically engineering the nervous system towards biomimetic control. However, a technology capable of directly manipulating the recruitment interface to enhance fatigue resistance remains challenging in the context of clinical efficacy and safety^24^. Indeed, if such translational challenges were resolved, such direct muscle augmentation technology could unlock promising new capabilities in muscle-actuated biohybrid systems.

An innervated skeletal muscle is not an ideal actuator for biohybrid systems, as it is volitionally controlled by the central nervous system (CNS), and due to its axonal biophysics, fatigues rapidly under continuous FES control^19,20^. Here we present a myoneural actuator (MNA) technology in a rodent model that fully redirects nervous system control to a computer-controlled modality, and augments fatigue resistance by manipulating axonal composition. By removing nervous system control through motor denervation, and reinnervating with a pure sensory nerve, we preserve muscle self-sustainability while enabling computer control via the restored neuromuscular junctions (NMJ) between the sensory nerve and muscle fibers using FES. Additionally, we show that the uniform axon size distribution of the reinnervated sensory nerve, compared to the native motor nerve, promotes enhanced fiber recruitment under FES. We endow the MNA with a closed-loop control architecture and reversible neural isolation from the CNS, providing external control authority over augmented actuation, without unwanted afferent feedback signals to the CNS (Fig. 1b). To demonstrate the potential applications of the MNA, we design a biohybrid neuroprosthetic interface and a biohybrid organ system, enabling modulation of neural afferents for bionic neurofeedback, and mechanical function of the small intestine, respectively. Our MNA technology offers design flexibility and controllability over neurally-isolated, fatigue-resistant actuation, while preserving innate muscle tissue properties, thus expanding the capabilities of implantable biohybrid systems.

## Results

### Myoneural actuator (MNA)

We sought to repurpose biological skeletal muscle as an actuator for biohybrid devices. To remove volitional control from the CNS, our design involves devervating the native motor nerve of the base muscle, which typically results in loss of contractility. To preserve muscle contractility, we reinnervate the MNA base with a pure sensory nerve^25,26^. We implemented the MNA in a rodent model, using the lateral gastrocnemius (LG) muscle as the actuator base and the sural nerve (SN) for sensory reinnervation (Fig. 2a and Extended Data Fig. 1).

**Fig. 2.**
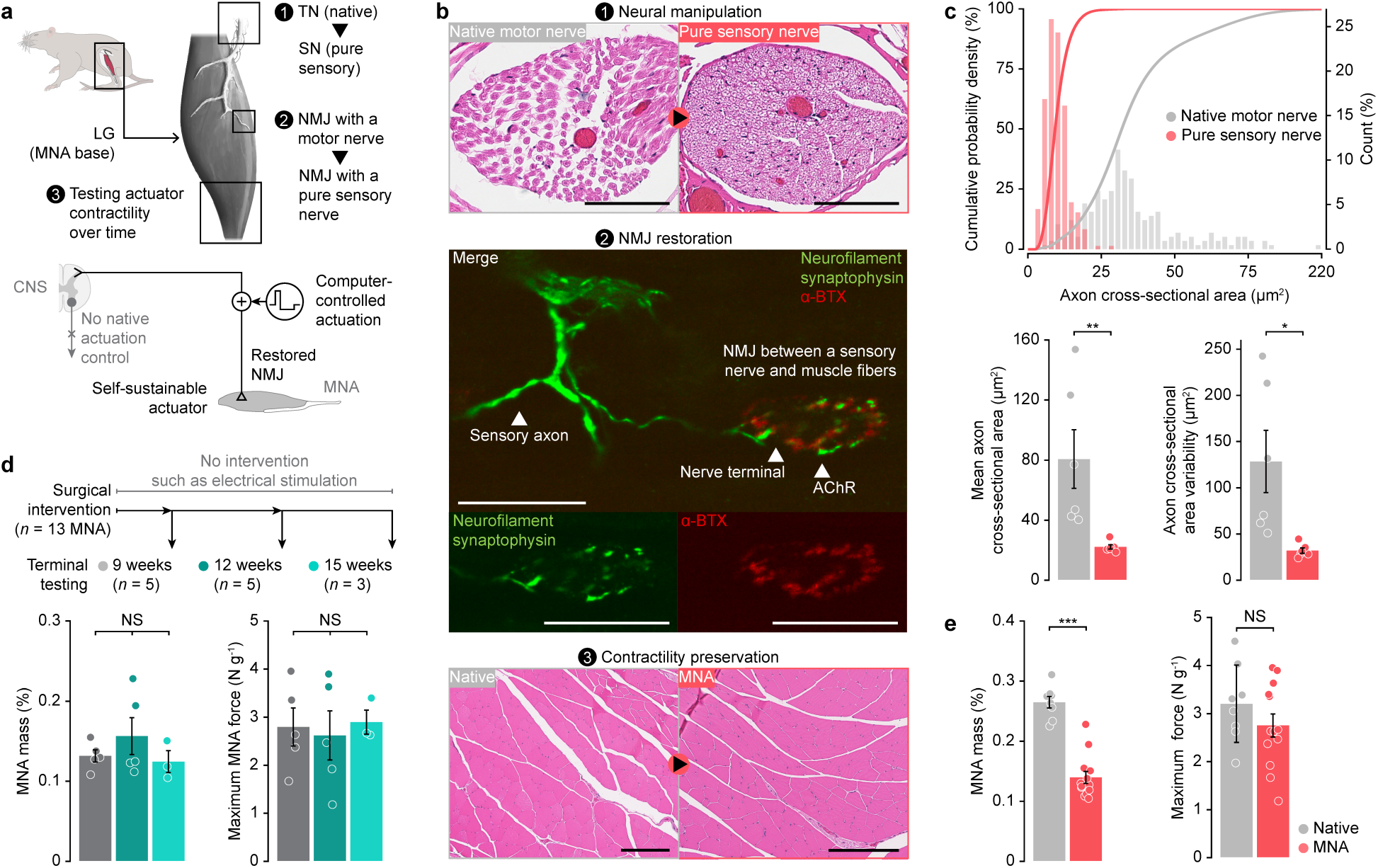
MNA provides a neural pathway for functional stimulation control and self-sustainability. **a**, MNA implementation in a rodent model and schematic of the redirected MNA neural pathway from the CNS to functional stimulation via computer control. LG, lateral gastrocnemius; TN, tibial nerve; SN, sural nerve; NMJ, neuromuscular junction. **b**, Representative histology (H&E) nerve cross-section of native motor nerve (TN) and pure sensory nerve (SN) used in the MNA. Scale bars, 50 μm (1). Representative restored neuromuscular junction (NMJ) between a pure sensory nerve and muscle fibers. Scale bar, 50 μm (2). Representative histology (H&E) of muscle fibers cross-section before and after myoneural manipulation. Scale bars, 200 μm (3). Extended Data Fig. 5 reports histological analyses in more detail. **c**, Axon cross-sectional area histogram, mean axon cross-sectional area and its variability for native motor nerves and pure sensory nerves used in the MNA (bars, mean ± SEM; *n* = 6 per cohort; area: two-sided Mann-Whitney U test, \*\**P* = 0.0022, variability: unpaired two-sided *t*-test, \**P* = 0.017). **d**, MNA mass and maximum force at various time points post-surgical intervention (bars, mean ± SEM; 9 and 12 weeks: *n* = 5, 15 weeks: *n* = 3; one-way analysis of variance (ANOVA), mass: *P* = 0.43, force: *P* = 0.91). NS, not significant. **e**, Mass and maximum force of native LG (Native) and MNA (bars, mean ± SEM; Native: *n* = 8, MNA: *n* = 13; mass: two-sided Mann-Whitney U test, \*\*\**P* = 2.5×10^-4^, force: unpaired two-sided *t*-test, *P* = 0.25).

Motor nerves, such as the tibial nerve (TN), exhibit variable axon diameters to supply distinct muscle fiber populations^19^. This non-uniform axonal composition contributes to the preferential recruitment of large-diameter fibers during FES, giving rise to unnatural motor unit recruitment and accelerating muscle fatigue^19,20^. In contrast, pure sensory nerves are known to have a more homogeneous axonal population^27,28^. To validate this, we conducted histological analyses of the TN that innervates the LG and the SN used for MNA reinnervation (Fig. 2b). As anticipated, the SN, a pure sensory nerve, exhibited smaller and more uniform axon cross-sectional area compared to the TN, a motor nerve (Fig. 2c). Thus, we hypothesized that the MNA reinnervated with a pure sensory nerve would demonstrate impartial fiber recruitment under FES control, enabling fatigue-resistant actuation while maintaining self-sustainability.

To test this hypothesis, we first conducted nerve-muscle histological analyses to investigate myoneural interactions resulting from reinnervation. Notably, reinnervation with a sensory nerve restored neuromuscular junctions (NMJs) (Fig. 2b), a phenomenon that, to our knowledge, has been elusive in the existing literature. This restoration enabled MNA modulation through the natural neural pathway via nerve stimulation, replicating the unique capabilities of a motor nerve while utilizing a pure sensory nerve for computer control. Histological analysis confirmed healthy muscle fibers in the MNA with no morphological difference or significant muscle fiber type transformation from native skeletal muscle (Fig. 2b, Extended Data Fig. 2).

### MNA sustainability without external interventions

Our MNA design eliminates efferent neural signaling from the CNS, resulting in muscle inactivity when the MNA is not under functional stimulation control. Prolonged inactivity could potentially reduce muscle mass and contractility, necessitating periodic functional stimulation interventions, such as FES, to maintain function^29,30^. To evaluate the long-term sustainability of the MNAs without such interventions, we measured the mass and maximum isometric force of three separate MNA cohorts at weeks 9, 12, and 15 post-implementation (Extended Data Fig. 3). No external interventions were applied until assessment. Our results revealed no significant differences in MNA mass or maximum isometric force across all time points (Fig. 2d). Furthermore, the MNA maximum isometric force was comparable to native muscles for a given mass (Fig. 2e). While there was an absolute loss of mass due to the reinnervation process, no additional atrophy was observed once reinnervation was complete (Fig. 2e). This opens the possibility for long-term enhanced recruitment of muscle fibers under FES control in a self-sustaining muscle actuator (Fig. 2a).

### Fatigue resistant MNA actuation

Histological analysis of the MNA revealed a more uniform axonal distribution in its reinnervating nerve compared to that of the native muscle. We hypothesized that this difference in axonal biophysics would translate to enhanced muscle fatigue resistance during repetitive and continuous actuation. To test this hypothesis, we evaluated the fatigue responses of the MNA and compared them to native LG muscle in two extreme scenarios: sequential twitch and continuous force production (Extended Data Fig. 3). Sequential twitch force responses were assessed over 450 cycles with 1-second resting intervals between single-pulse stimulations (Fig. 3a). As anticipated, the MNA cohort demonstrated superior preservation of output force during sequential single-pulse actuation compared to the native cohort (Fig. 3a and 3b). Furthermore, the MNA cohort reached an earlier equilibrium between fatigue and recovery during single-pulse stimulations, whereas the native cohort experienced continuous fatigue (Fig. 3b). As a result, the MNA exhibited a significantly lower twitch force loss rate compared to the native cohort (Fig. 3c) and showed more stable actuation with reduced force variability (Fig. 3d). Under continuous actuation without resting periods, the MNA cohort demonstrated a 260% improvement in fatigue resistance compared to the native cohort (Native: 5.19 ± 1.16 s, MNA: 18.67 ± 2.96 s) (Fig. 3e).

**Fig. 3.**
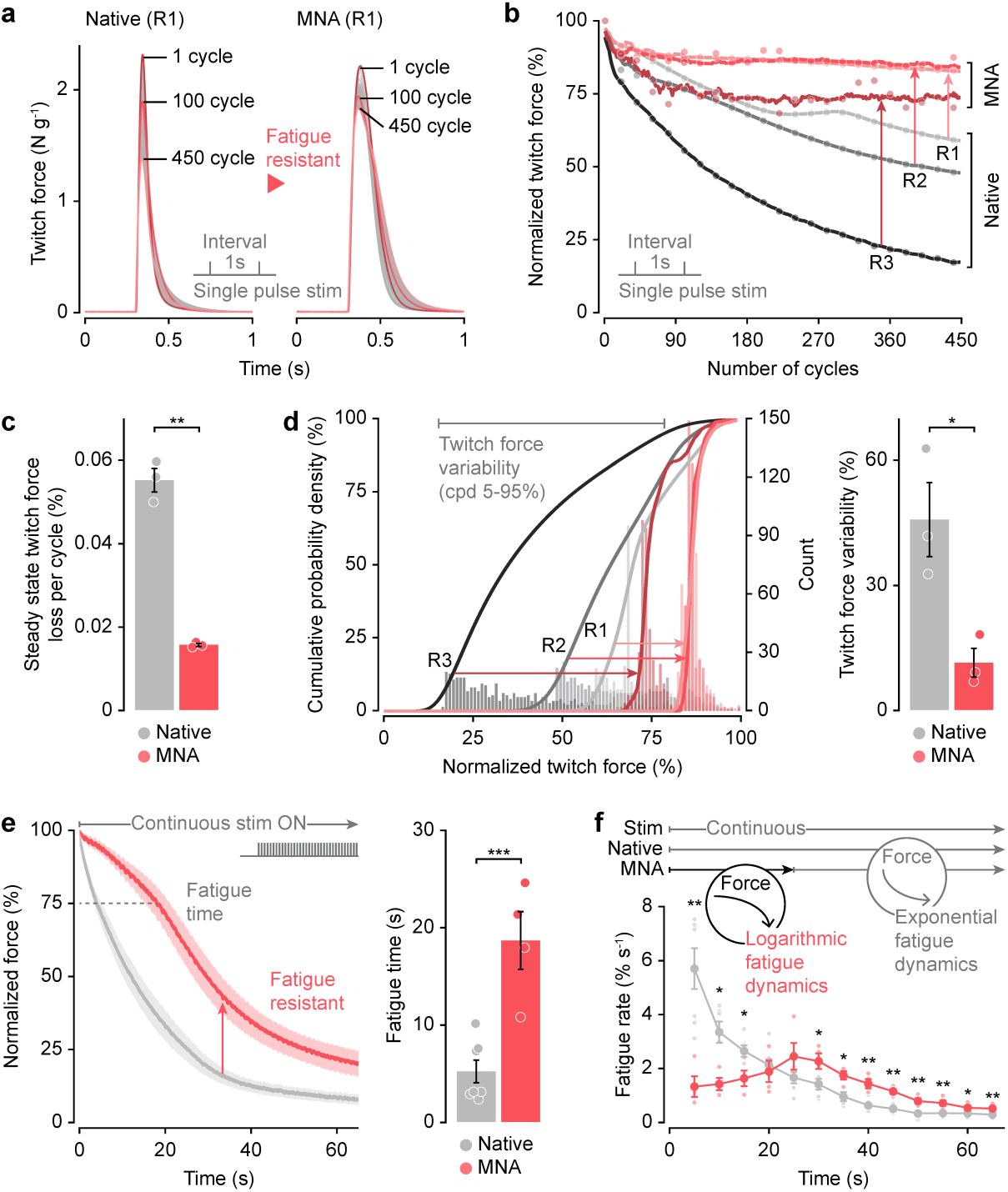
MNA enables fatigue resistant actuation. **a**, Representative sequential twitch force responses for a native muscle and MNA (*n* = 450 cycles per muscle). R1, Rat1. **b**, Summary of sequential twitch force responses for both cohorts (*n* = 450 cycles per muscle). R1-R3, rat1-rat3. **c**, Steady state twitch force loss of native and MNA between cycles (bars, mean ± SEM; *n* = 3 per cohort; paired two-tailed *t*-test, \*\**P* = 0.0061). **d**, Twitch force response histogram, cumulative probability density (cpd), and variability for both cohorts (bars, mean ± SEM; *n* = 3 per cohort; paired two-tailed *t*-test, \**P* = 0.030). **e**, Force responses under continuous stimulation for both cohorts (lines and bars, mean ± SEM; Native: *n* = 7, MNA: *n* = 4; unpaired two-tailed *t*-test, \*\*\**P* = 0.00067). **f**, Fundamentally different fatigue resistant muscle dynamics of the MNA cohort compared to those of the native cohort (bars, mean ± SEM; Native; *n* = 7, MNA: *n* = 4; Mann-Whitney U-tests, \**P* < 0.042, \*\**P =* 0.0061).

We investigated whether this augmented fatigue resistance stemmed from simply having a longer fatigue time while sharing the same exponential fatigue dynamics of native muscles (Fig. 3f). Unexpectedly, we found fundamentally different fatigue muscle dynamics in the MNA cohort compared to the native cohort, characterized by additional logarithmic fatigue dynamics (fatigue resistant) compared to the common exponential fatigue dynamics. This fatigue resistant dynamics enabled the MNAs to sustain high forces more effectively under continuous actuation compared to native muscles.

### MNA extended closed-loop control

We hypothesized that the fatigue-resistant muscle actuation provided by the MNA would enable extended force controllability compared to native muscles, even when implemented in a closed-loop control architecture designed to compensante for force tracking errors. To test this hypothesis, a closed-loop force control architecture with a fixed feedback gain was implemented for both cohorts, with target force values set at 30%, 50%, and 70% of maximum force (Fig. 4a, Extended Data Fig. 3). We conducted 45 sequential cycles for each target value to assess control performance (Fig. 4b). Initially, the closed-loop controller effectively tracked each target value for both the native muscle and the MNA (Fig. 4c and 4d). However, as the number of cycles increased, the native muscle rapidly lost force controllability due to fatigue, regardless of the target values (Fig. 4c and 4d). In contrast, the MNA maintained force controllability across varying target levels, even as the number of cycles increased (Fig. 4c and 4d). Our findings highlight that fatigue in native muscles under FES control remains a critical limiting factor in actuation, despite compensatory efforts of a closed-loop controller. The fatigue resistant actuation enabled by our MNA technology improves the fundamental capability of muscle-actuated systems by preserving force controllability over extended actuation periods.

**Fig. 4.**
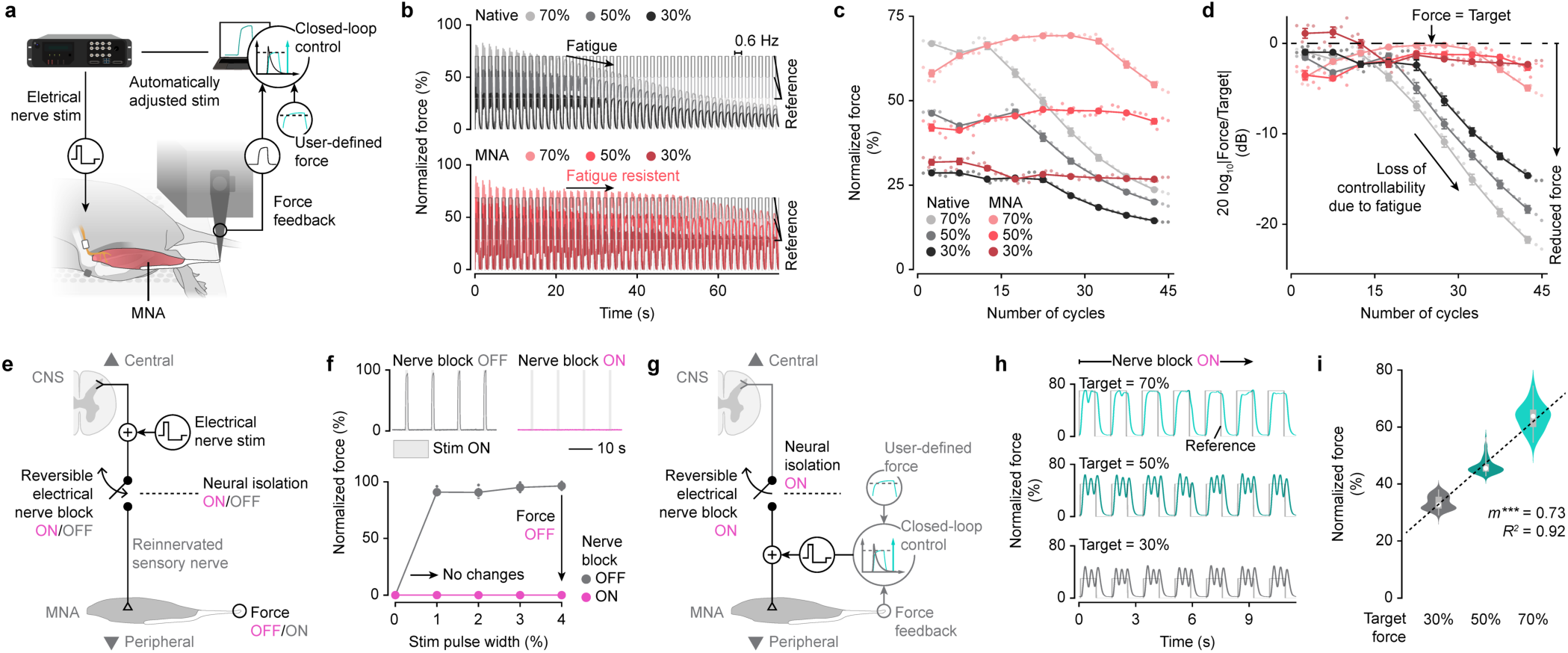
MNA enables extended closed-loop control with neural isolation from the CNS. **a**, Experimental setup for closed-loop force control. **b**, Closed-loop control performance for the native muscle and MNA across various force target values. **c** and **d**, Summary of closed-loop control force outputs (c) and the gain between target and output values (d) (*n* = 45 cycles per each target value; large dots, means for every *n* = 5 cycles). **e** and **f**, To verify nerve block effect, reversible electrical nerve block was applied distal to the nerve stimulation site for MNA modulation (e). Nerve block effectively shunted neural conduction, silencing the MNA force output in response to nerve stimulation (f). **g**-**i**, To demonstrate MNA closed-loop controllability with neural isolation, nerve block was applied proximal to the nerve stimulation site to neurally isolate the MNA from the CNS during actuation (g). The MNA demonstrated the capability to modulate force at varying levels under nerve block (h and i) (30%: *n* = 89 cycles, 50%: *n* = 85 cycles, 70%: *n* = 33 cycles; large white dots, medians; boxes and whiskers, interquartile ranges and adjacent values). Slopes (*m*) and *R*^2^ between target and output force values are reported (\*\*\**P* < 0.001).

### MNA modulation with reversible neural isolation from the CNS

Modulating the MNA through nerve stimulation inevitably activates neural pathways towards both the distal (MNA) and the proximal (CNS) end. This neural activation can induce unintended sensations, pain, or disrupt natural neural signaling crucial for neuromotor control if transmitted to the CNS. Therefore, isolating the proximal signaling pathway during MNA modulation is essential to enable actuation without potentially provoking adverse neural activation and broaden the applicable stimulation parameters and protocols. To achieve this, we integrated reversible electrical nerve block to our MNA technology^31^.

We first verified the efficacy of the nerve block by applying it distally to the nerve stimulation for MNA modulation, positioned between the nerve stimulator and the MNA (Fig. 4e). Activation of the nerve block effectively shunted the distal neural pathway, silencing the MNA response to nerve stimulation (Fig. 4f), confirming the effectiveness of the nerve block in isolating the neural pathway at the point of application. Subsequently, we conducted closed-loop control with the nerve block applied proximally to the nerve stimulation, positioned between the nerve stimulator and the CNS (Fig. 4g). This setup allowed us to assess MNA modulation while being neurally isolated from the CNS. Our results demonstrated that the MNA could achieve varying levels of target force values while under nerve block (Fig. 4h), with high linearity observed between the target and output force (Fig. 4i). These findings validated the ability of the MNA technology to provide muscle actuation while being neurally isolated from the CNS during operation.

### Biohybrid neuroprosthetic interface for neural feedback of a bionic limb

Muscle-tendon proprioception, the sensation of muscle-tendon length, velocity and force, relies on the activation of mechanoreceptors within muscle-tendon constructs (mechanoneural actuation)^32^. The ability to control mechanoneural actuation presents an opportunity to modulate proprioception, which has diverse applications including providing neural feedback for bionic limbs or virtual avatars in the metaverse (Fig. 1a).

Here we demonstrate the application of our MNA technology within the context of a biohybrid neuroprosthetic interface for bionic integration after limb amputation. We conceived the Proprioceptive Mechanoneural Interface (PMI) to both provide efferent control over a prosthesis, as well as to modulate proprioceptive neural afferents in the amputated residuum in response to external bionic joint states and torques (Fig. 5a). The PMI consists of a MNA linked in series to a residual muscle end organ, sensing of the end organ’s state and force, and a closed-loop control system wherein the length, speed and force of the end organ can be directly modulated to control neural afferents (Fig. 5a and 5b). In principle, multiple PMIs can be constructed for each residual muscle to fully mimic intact muscle dynamics in response to bionic joint positions, movements and torques. At least two PMIs are required for a single joint, one PMI for a flexor and another for an extensor. In this framework, the afferent feedback of prosthetic states and torques is achieved by directly modulating the states and forces of the end organs through MNA closed-loop control. In addition, the measured states and forces of the muscle end organs of the PMIs under CNS control, serve as efferent neural signals for prosthetic control, creating a bidirectional neuroprosthetic interface (Fig. 5a).

**Fig. 5.**
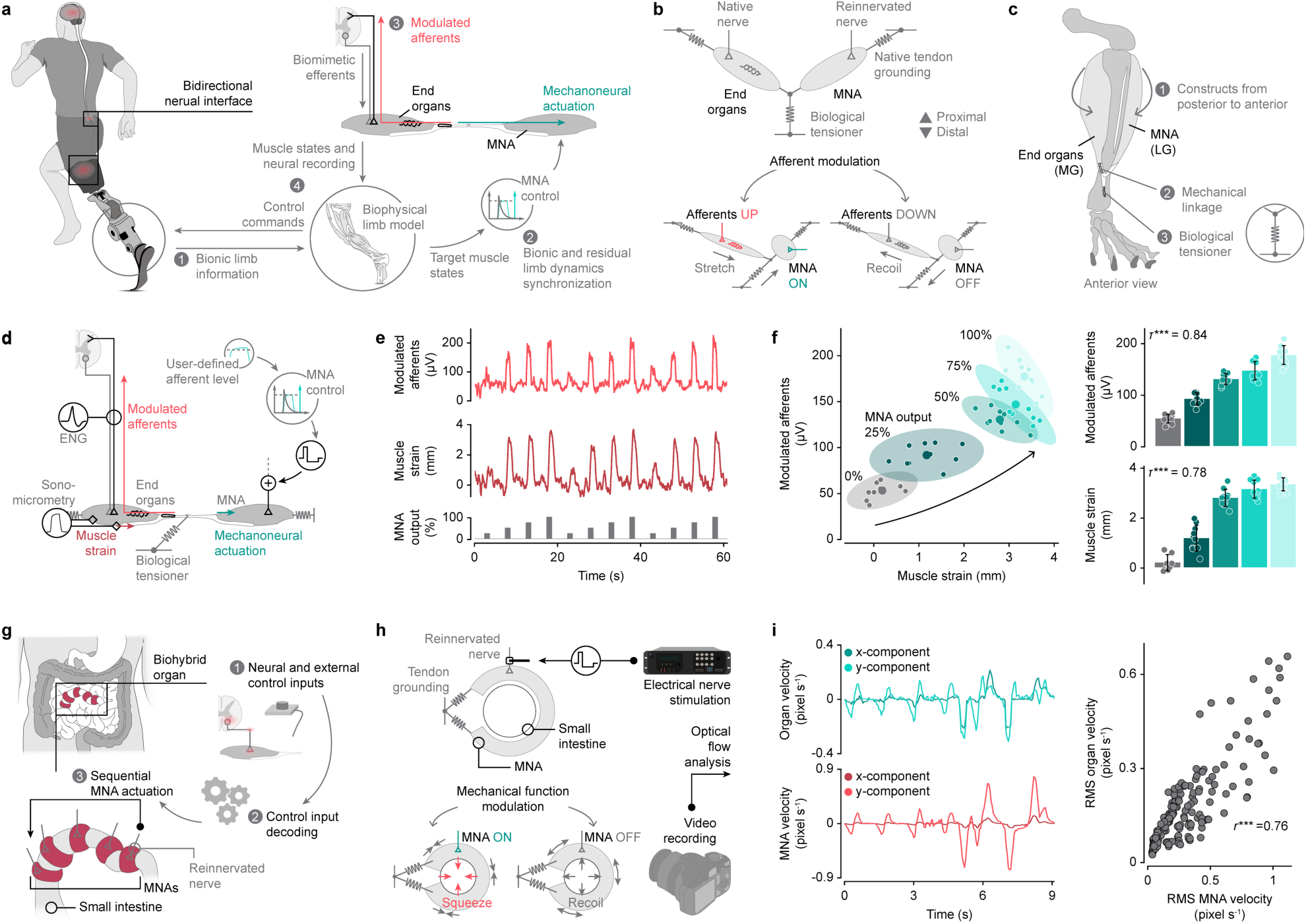
Biohybrid neuroprosthetic interface and biohybrid organ system using MNAs. **a**, Conceptual diagram of a biohybrid neuroprosthetic interface, the Proprioceptive Mechanoneural Interface (PMI). The PMI consists of a residual muscle with state and force sensing (end organ) serially coupled with a MNA, enabling both a closed-loop mechanoneural control of the end organ to provide proprioceptive feedback from a bionic limb, and an efferent signal for neural control of the bionic limb. **b**, Schematic of the PMI functional mechanism design. **c**, Surgical design of the PMI in a rodent model, using the medial gastrocnemius (MG) to mimic a residual muscle in an amputation scenario, with the MNA constructed from the LG. **d**, Schematic of the experimental setup for PMI evaluation. End organs strain and modulated afferents were assessed through sonomicrometry and electroneurography (ENG), respectively. **e** and **f**, The PMI demonstrated the ability to modulate end organ strains and neural afferents at different levels through MNA actuation (e). As MNA output increased, both muscle strains and modulated afferents increased (*n* = 10 cycles per each level) (f). Kendall’s tau coefficients (τ) between MNA outputs, residual muscle strains, and modulated afferents are reported (afferents: \*\*\**P* = 1.9 ×10^-15^, strains: \*\*\**P* = 1.8 ×10^-13^). **g**, Conceptual diagram of a biohybrid organ system designed for the small intestine. Neural signals from the nervous system, along with external signals, can serve as control inputs for the biohybrid organ system. **h**, Schematic of the biohybrid organ system functional design and the experimental setup for evaluation. **i**, Optical flow analysis revealed a strong correlation in movement between the MNA and the small intestine. Pearson correlation coefficient (*r*) between root-mean-square (RMS) velocities of organ and MNA is reported (*n* = 186, \*\*\**P* = 2.3 ×10^-36^).

In our rodent model demonstration, we utilized the medial gastrocnemius (MG) muscle to simulate an end organ in an amputation scenario. The MG was surgically coupled to the MNA at the anterior tibia site, with the distal MG tendon serving as a biological linear tensioner (Fig. 5c and Extended Data Fig. 4). We evaluated end organ strain using sonomicrometry and modulated afferents with electroneurography (ENG) recordings (Fig. 5d). By incrementally increasing MNA actuation levels, we tested the PMI capability to modulate afferents from the end organs (Fig. 5e). Our findings revealed that both end organ strains and modulated afferents increased with higher MNA actuation levels (Fig. 5f). This highlights the integration of the MNA technology with biological sensory organs for neural signal modulation through mechanoneural actuation.

### Biohybrid organ system for mechanical modulation

Various organs facilitate vital functions through mechanical movements. For instance, the contraction of the thoracic diaphragm facilitates respiration^8^, while the contraction of the bladder, small intestine, and vascular system enables excretion, digestion, and hemodynamic control, respectively^33–35^. Furthermore, these organs have a direct influence on our behaviors and mental states, such as reward and satiation^36–38^. The capacity to control organ mechanics holds the potential to develop technologies that can address organ failure, physiological conditions, and even provide physiological feedback in virtual environments like the metaverse.

Here we demonstrated the application of our MNA technology as a mechanical emulator of organ function. We designed a biohybrid small intestine, wherein the MNA wraps around the organ to mimic its mechanical squeezing function (Fig. 5g, Extended Data Fig. 5). This biohybrid organ system can be controlled using neural signals from the nervous system, along with external signals. Multiple MNAs can be employed to emulate sequential actuation such as peristalsis. In our demonstration, we implemented the biohybrid small intestine in a rodent model using a single MNA (Fig. 5h). Optical flow analysis revealed synchronized movements between the MNA and the small intestine at varying actuation levels (Fig. 5i), validating its capability to modulate organ mechanics.

## Discussion

A fully biointegrable actuator with high-performance, continuous actuation could enable modulation of a myriad of biological functions, from restoring organ function, to generating neural afferent feedback for neuroprostheses and virtual experiences. Despite substantial improvements in synthetic actuator and biofabrication technologies, an actuator with these features does not yet exist. In this study, we introduce the MNA, an engineered regenerative construct with the bioimplantable and actuation properties of skeletal muscles, while showing fatigue-resistant continuous actuation under computer control.

### MNA mechanism

The MNA is based on a myoneural framework that transforms a native skeletal muscle into a fatigue-resistant actuator under FES computer control. The actuator is constructed by redirecting native CNS motor control through denervation of its native motor nerve, followed by reinnervation using a sensory nerve to enhance fatigue resistance. We provide morphological and electrophysiological evidence that the sensory nerve reinnervates muscle fibers. This enables the preservation of muscle function without external interventions and allows muscle fibers to be controlled via the natural neural pathway. The MNA exhibited significantly enhanced fatigue-resistance in both open-loop and closed-loop settings, along with fundamentally-distinct, fatigue dynamics compared to native muscles without the myoneural manipulation.

The MNA construction involves reinnervation by a pure sensory nerve, guided by a twofold design rationale: to preserve muscle function, as supported by sensory protection interventions^25,26^, and to modify the recruitment biophysics under electrical nerve stimulation. During extraneural stimulation of peripheral nerves with varying axon sizes, such as a motor nerve, the lower resistance of large-diameter axons leads to preferential activation of motor units that typically innervate fast-twitch muscle fibers, thereby accelerating muscle fatigue^19,20^. We thus surmise that the more uniform fiber-size distribution of sensory nerves, compared to motor nerves, underpins the enhanced fatigue resistance observed in the MNA, providing a potential solution to this persistent limitation. This is in line with recent evidence that suggests that by uniformly expressing light-sensitive ion channels in nerves, muscles can be optically controlled in an orderly natural manner^19^ with improved recruitment profiles, controllability and fatigue resistance^20^. This improved muscle modulation arises from the greater access to axons in the nerve, allowing the activation of most fibers and not only large ones.

Seminal studies on cross-reinnervation have elucidated muscle fiber type transformations^39,40^, while recent work has shown that peripheral motor nerve transfers induce a donor-specific shift to slow fiber types, potentially increasing fatigue resistance^41^. However, these transfers also result in higher conduction velocities, suggesting that motor units contain large-diameter fibers, which are more prone to fatigue. These studies suggests that our sensory nerve transfer could induce fiber type transformation, however, we did not observe a significant shift (Supplementary Data Fig. 2). Removing motor control may prevent fiber transformation, as shown in studies where the absence of spinal cord signals^39,40^ or absence of changes in the pattern of impulse activity reaching the muscle had no significant effect on fiber transformation^40^. Our prevailing hypothesis is that the observed fatigue-resistance stems from the axonal biophysics of the sensory nerve; however, we cannot fully rule out the possibility of fiber type transformation, which could have been revealed with a different immunohistological procedure^42^.

### Biohybrid system applications

The MNA technology enables a range of biohybrid applications in medicine, including the neuroprosthetic interface and the organ system presented here. We present a new mechanoneural interface, the PMI, for neural efferent-afferent modulation of prosthetic limbs^43^. In contrast to the agonist-antagonist myoneural interface (AMI)^44–47^, the PMI possess several advantages^43^. Firstly, the PMI can provide independent force afferent feedback through its agonist muscle without activating the antagonist muscle, enabling afferent feedback of external forces, such as gravity and inertia felt by the prosthesis, on the agonist muscle end organ without unwanted antagonist activation (or vice versa). Secondly, in contrast to the AMI, wherein biological transmission is physically determined by linking muscles together, the PMI allows for accurate biological transmission emulation in software based on an intact biophysical model. Finally, for proximal amputations, the regenerative AMI requires a two-stage surgery^17^. Because the PMI pairs are not physically attached, the regenerative PMI can be done in a single-stage surgery. The PMI is part of a growing number of mechanoneural constructs that aim to provide improved neural control and natural afferents. A number of these interfaces, such as the cutaneous mechanoneural interface^18^ (CMI), the composite regenerative peripheral nerve interface (C-RPNI), the dermal sensory RPNI (DS-RPNI), targeted sensory reinnervation (TSR), and the sensorimotor interface^17,48^, could benefit from a bioimplantable fatigue-resistant actuator like the MNA.

We also introduced a biohybrid organ system to demonstrate how organ mechanics can be modulated with the MNA. Conditions such as ileus, diabetic enteropathy, and Crohn’s Disease can impair intestinal contractions, hindering movement of food and nutrients through the GI tract^49–51^. We showed how the MNA can interface with the small intestine to modulate its mechanics on demand, laying the foundation for a biohybrid organ system to restore GI tract function. Furthermore, mechanical actuation of the small intestine and the GI tract can trigger a cascade of physiological effects, including gene expression from mechanosensors activation, hormone release such as GLP-1, and modulation of neurotransmitters and enzymes^52^. These changes can lead to improved nutrient absorption and energy balance, gut-brain axis modulation, and influence neurological functions such as mood, anxiety and cognitive function^37,38^. The MNA could be used as a tool to modulate organ mechanics and to study their downstream effects.

Due to the MNA neural isolation from the CNS, it is an attractive tool for autonomic function control. Beyond modulating neural sensory afferents and the GI tract, autonomic functions such as respiration and urinary control could be achieved through mechanical modulation of the lungs and bladder. For respiratory modulation of individuals with ventilatory insufficiency, a set of MNAs could be coupled to the diaphragm and intercostal muscles, facilitating coordinated contraction during inspiration, while respiratory sensor signals inform the timing and intensity of the actuation (Extended Data Fig. 6). For urinary modulation, one or more MNAs could be wrapped around the detrusor muscle of the bladder to generate compression, assisting in urinary excretion through control inputs, such as smartphone commands (Extended Data Fig. 6).

While biohybrid system applications are promising, understanding tissue interactions between the MNA and distinct tissues, such as smooth or cardiac muscle, is critical. Insights from dynamic cardiomyoplasty, where a skeletal muscle is wrapped around the heart to assist pumping, revealed that differences in electrical conduction properties between skeletal and cardiac muscle can lead to ventricular arrythmias and fibrillation^53^. By engineering the tissue interface to electrically isolate the MNA from the myocardium, the MNA could help modulate heart mechanics. Unlike dynamic cardiomyoplasty, where native skeletal muscle requires extensive training to become fatigue-resistant^53^, the MNA could offer fatigue resistance without such conditioning. Additionally, the muscle used in cardiomyoplasty retains its original motor nerve, limiting neural isolation from the CNS and hindering computer control. Nerve stimulation to the MNA could be synchronized with ventricular systole for closed-loop assistance (Extended Data Fig. 6). A computational framework based on the functional demands and target tissue properties, would prove useful to come up with the adequate design choices for the MNA for particular biohybrid applications, such as size, tissue coupling modality, and stimulation parameters (Extended Data Fig. 6).

### Translation

Because the MNA is based on a reconstructive surgical technique to engineer skeletal muscle and standard nerve cuff electrodes, it is readily translatable for a number of medical applications. Similar surgical techniques are commonplace in facial nerve and brachial plexus surgeries, as well as in reconstructive approaches for bionic integration^17,54^. During the same surgery, a wireless pulse generator and nerve electrodes, which are commonplace, can be implanted. Additionally, minimally-invasive tissue sensing techniques would provide end organ state and enable wireless closed-loop MNA control (Extended Data Fig. 6)^55,56^. We showed that the MNA does not require external interventions for reinnervation and long-term sustainability. The sustainability data presented here is the most extreme case, external interventions such as stimulation protocols through the implanted stimulator could increase MNA sustainability and capacity, as well as transform muscle phenotype for desired functionality^30,57^. Similarly, the same stimulator can be used for reversible electrical nerve block to prevent unintended signaling during operation^31^. Nerve block would only be required if the stimulation parameters required for MNA control cause undesired or painful sensations. The relationship between stimulation parameters and evoked sensations could be investigated in patients with sensory protection to evaluate if nerve block is necessary^25,26^.

The use of an autograft skeletal muscle avoids problems of infection, rejection, and immunosuppression that will be encountered with allografts, xenografts or engineered tissues^58,59^. Additionally, the MNA procedure would be relatively simple and safe compared to implantable mechanical assist devices or organ transplants^8,60,61^. During reconstructive amputation procedures, skeletal muscle tissue from the same limb that otherwise would be discarded becomes available to create constructs like the MNA^17^. In contrast to procedures like cardiomyoplasty, where the muscle needs to be in close proximity to the heart due to the requirement of an intact neurovascular bundle^53^, in principle the MNAs can be created from skeletal muscles from any location. Muscles such as latissimus dorsi from the back, rectus abdominis from the abdomen, or pectoralis major from the chest, are commonly used as autografts in other procedures^62–64^. Sensory nerves that innervate skin areas in proximity to the organ of interest can be used for constructing the MNA. Considerations to select MNA components such as muscle and nerve size are important to meet functional clinical demands (Extended Data Fig. 6).

In summary, we have shown that the MNA is a biointegrable regenerative tissue construct with augmented fatigue resistance, capable of continuous actuation under computer control. The design approach to engineer biology through regenerative techniques, alongside with clinically-approved stimulation and sensing technology, enables readily clinical translation for a number of important medical conditions. The MNA technology expands the fundamental capabilities of muscle-actuated, implantable biohybrid systems.

## Methods

### Animal husbandry

All surgical and experimental procedures are approved by the Committee on Animal Care (CAC; protocol 2203000299A007) at the Massachusetts Institute of Technology (MIT) and conducted in accordance with ethical guidelines. The study involved a total of *n* = 30 female Lewis rats (273.6 ± 7.1g body weight). The rats were kept in pairs under a light/dark cycle of 12 hours.

### Myoneural actuator (MNA) design and surgical implementation

This study aimed to create an MNA from a native muscle (MNA base) by redirecting motor control from the CNS to computer control while preserving self-sustainability. To achieve this, we reinnervated the MNA base with a pure sensory nerve and subsequently denervated the native motor nerve, preventing the loss of muscle contractility typically caused by denervation. We hypothesized that neuromuscular junctions (NMJs) between the sensory nerve and muscle fibers would reform, enabling the neural pathway to support computer control. Furthermore, due to the more uniform axon size distribution in sensory nerves compared to motor nerves, we hypothesized that sensory-nerve activation would recruit a broader range of muscle fibers more evenly under electrical stimulation, resulting in fatigue-resistant actuation. The surgical procedures for MNA implementation in a rodent model are shown in Extended Data Fig. 1. The lateral gastrocnemius muscle (LG) served as the MNA base, reinnervated by the sural nerve (SN, pure sensory nerve). A 1.5 cm incision was made on the lateral hind limb and biceps femoris (BF) to expose the tibial nerve (TN, native motor nerve) and SN. After denervating the TN branch to the LG, the proximal TN was sutured to the BF to prevent unintended reinnervation. The distal SN was transected, embedded into a muscle pocket on the LG, and the pocket was sutured closed. All incisions were closed with sutures. Surgeries were conducted under aseptic conditions and general anesthesia (2% isoflurane).

### Histological analysis

To verify MNA mechanisms, we conducted histological analysis on the native LG (without myoneural manipulation), MNAs, and their associated nerves (TN and SN). Tissues were carefully dissected and fixed in 4% PFA via immersion overnight at 4°C. Tissues were then washed three times with 1X PBS (10 minutes each). For NMJ and fiber type analysis, tissues were then cryoprotected in 30% sucrose for 48 hours at 4°C. Then, tissues were embedded in OCT and snap-frozen with liquid nitrogen and dry ice. For NMJ analysis, muscles with their nerve were sectioned longitudinally at 20 μm on a cryostat, mounted on glass slides, and stored at -20°C overnight. For fiber type analysis, muscles were sectioned cross-sectionally at 10 μm on a cryostat, mounted on glass slides, and stored at -20°C overnight. Afterward, sections were treated with 0.5% PBST for 15 minutes at room temperature. Blocking was performed with a solution containing 5% normal donkey serum, 0.1% Triton X-100, 3% bovine serum albumin, and 0.09% sodium azide in 1x PBS for 60 minutes at room temperature. Slides were then washed three times with PBS for 5 minutes. NMJ slides were incubated with primary antibodies anti-neurofilament-L at 1:1000 (Cell signaling catalog #2837) and anti-synaptophysin at 1:1000 (Cell signaling catalog #36406) overnight at 4°C. Sequential muscles slides for fiber type analysis were incubated with primary antibodies anti-myosin fast clone MY-32 at 1:500 (Sigma-Aldrich catalog #M1570) and anti-myosin slow clone NOQ7.5.4D at 1:10000 (Sigma Aldrich catalog #M8421) overnight at 4°C. Then, slides were washed with washing buffer three times for 10 minutes. NMJ slides were incubated with secondary antibody Alexa Fluor 488 at 1:1000 (abcam catalog #ab150077) and Alpha-Bungarotoxin conjugated with Alexa Fluor 594 at 1:1000 (Invitrogen catalog #B13423) for 2 hours at room temperature. Muscle slides were incubated with secondary antibody Alexa Fluor 488 at 1:1000 (Invitrogen catalog #A-21121) for 2 hours at room temperature. Slides were then washed with washing buffer three times for 10 minutes, excess liquid was removed, mounting media was added, and cover slips were applied and secured. For axon cross-sectional area and muscle fiber morphology analysis, after fixing and washing, tissues were stored in 75% ethanol followed by paraffin processing. Nerve and muscle tissues were sectioned cross-sectionally at 5 μm. Sections were then stained with haematoxylin and eosin. Images were acquired using a high-resolution multi-channel confocal system on a Nikon Eclipse Ti-E inverted microscope body with a Zyla 5.5 camera and an Andor spinning disk (SCU-W1 Tokogawa), and a Leica AT2 slide scanner. A 10X, 20X air objectives and a 40X water objective were used. A calibration slide was sued to determine image scale bars. Nikon element software version 4.0 and CellPose were used for processing data^65,66^. Axon cross-sectional area was computed using segmented images from Cellpose Cyto3 model. After segmentation, the model output included the area in pixel of each segmented region or interest (ROI). From this, the cross-sectional area was determined by converting the ROI pixel count into square micrometers using the scalebar as reference. Fiber type analysis was also performed using segmented images from Cellpose Cyto3 model. Images were converted to grayscale before segmentation. From the output, the total number of pixels from all ROIs was then summed to obtain the total pixel area of the fluorescent muscle fibers. The total muscle area was found by counting the total number of pixels in the entire region. The percentage of fluorescent muscle fibers for each image was then found by dividing the total fluorescent area by the total muscle area (Supplementary Data Fig. 2).

### Long-term MNA self-sustainability

MNA self-sustainability was evaluated by measuring maximum isometric force and mass at 9, 12, and 15 weeks, with additional native LG for baseline evaluation. To ensure self-sustainability without external intervention, separate MNA cohorts were prepared for each time point, and no stimulation was applied until assessment. Force measurements were conducted using a custom experimental apparatus comprising an adjustable heated bed and a force transducer (305C-LR, Aurora Scientific, USA)^20^ (ref: closed-loop optogenetic). The procedures for force measurement are illustrated in Extended Data Fig. 3. Approximately 4 cm lateral tibial incision exposed the MNA or native LG by retracting the biceps femoris (BF). The sciatic nerve branches to surrounding muscles such as tibial anterior or soleus were transected to prevent unintended activation from potential current leakage during electrical stimulation. SN or TN surrounding tissues were cleared for electrode placement, and the Achilles tendon was transected as distally as possible to isolate the MNA or LG from the medial gastrocnemius (MG). The animal was then positioned in the apparatus, with the patellar tendon and anterior talofibular ligament sutured to the heated bed for stabilization, and the distal tendon sutured to the force transducer. Muscle tension was adjusted to position the muscle-tendon unit at its resting length. Electrical stimulation was applied through a hook electrode (SN for MNAs, TN for native LG) using a 1-second stimulation (70 μA, 40 Hz) with incremental pulse widths (PW). PW increased by 0.05% up to 0.25%, then by 0.25% to 1%, and by 1% thereafter until force plateaued. Following characterization, MNA or LG mass was recorded and normalized to the pre-test body weight. Maximum isometric force was normalized to the weight of each actuator.

### Fatigue response and dynamics

MNA fatigue responses during single-pulse and continuous stimulation were evaluated to assess potential fatigue-resistant actuation compared to native LG. Separate cohorts were prepared for each stimulation protocol to prevent carryover effects. Experimental preparation followed the ‘long-term MNA self-sustainability’ protocol. For single-pulse testing, 450 pulses (70 μA, 2 ms PW) were delivered sequentially with 1-second intervals. For continuous stimulation, 70 μA, 40 Hz, and 4% PW were applied for 5 minutes. Force data were normalized to maximum values. Fatigue resistance during single-pulse stimulation was assessed by calculating the steady-state twitch force loss per cycle, determined by the average force loss rate over the last 100 cycles (from cycles 346-350 and 446-450). Force consistency was evaluated by the 5-95% cumulative probability density bandwidth (twitch force variability). For continuous stimulation, fatigue resistance was quantified by the time it took for the output force to decrease to 75% of its maximum (fatigue time). Differences in continuous fatigue dynamics between MNA and native LG were analyzed by comparing fatigue rates at each time point, calculated from average force values every 5 seconds.

### Fatigue resistant closed-loop force control

Closed-loop force control was implemented using the same experimental setup to assess the impact of fatigue resistance on force controllability. Target force values were set at 30%, 50%, and 70% of the maximum output force for each MNA and native LG. A sequence of 45 cycles at 0.6 Hz were recorded for each target level. Closed-loop controllability was evaluated based on the tracking performance and the gain between output force and target values, calculated as 20log_10_(Force/Target).

### Closed-loop control with neural isolation

Electrical stimulation propagates to both the distal MNA and proximal CNS. To achieve neural isolation and prevent unintended signaling to the CNS during MNA operation, a reversible electrical nerve block was integrated. High-frequency electrical stimulation (8 V peak-to-peak, 20 kHz) was applied to block neural signals. Validation of the nerve block via electroneurography (ENG) recording was challenging due to stimulation artifacts; therefore, verification was based on muscle force output, consistent with previous studies. Two nerve cuffs were placed on the SN of the MNA: nerve block was applied through the distal cuff, while the proximal cuff attempted to activate the MNA. Nerve block performance was tested at varying stimulation levels (1%, 2%, 3%, 4% PW) by recording output force. Baseline force was collected without stimulation. Ten cycles of force output per condition were recorded using the same experimental apparatus as prior tests. For closed-loop force control with neural isolation, nerve block was applied via the proximal cuff while the distal cuff controlled the MNA. The protocol followed the previously described closed-loop force control method but with a different number of cycles to account for enhanced MNA fatigue resistance (30% target: *n* = 89 cycles; 50%: *n* = 85 cycles; 70%: *n* = 33 cycles).

### Proprioceptive Mechanoneural Interface (PMI)

To demonstrate the potential of MNA technology, we developed a novel neuroprosthetic interface, the Proprioceptive Mechanoneural Interface (PMI), enabling neural afferent modulation from a residual muscle for neuroprosthetic limbs. The PMI comprised a residual muscle in an amputated limb, serially coupled with an MNA. This setup allowed the MNA to modulate the stretch of the residual muscle-tendon complex, controlling mechanotransduction in sensory end organs to generate proprioceptive feedback for neuroprosthetic limb. Procedures for PMI implementation in a rodent model are illustrated in Extended Data Fig. 4. The MG was used to simulate a residual muscle and was serially coupled with an MNA. SN access and MNA separation from MG followed prior procedures, but the tibial nerve branch to MG remained intact for afferent recording. The residual muscle and MNA were repositioned to the anterior tibia and surgically coupled via one tendon, with the other tendon secured to distal tissue near the ankle joint to prevent slippage. To assess mechanoneural actuation and afferent modulation, sonomicrometry crystals and a nerve cuff were placed on the MG and its tibial nerve branch. Muscle strain was recorded at 200 Hz, and raw ENG at 6 kHz. The raw ENG underwent band-pass filtering (second-order Butterworth, pass-band: 300–3000 Hz), artifact blanking (2 ms post-stimulation) based on stimulation triggers, rectification-bin integration (RBI) in a 4 ms window, and smoothing (250 ms moving average). A hook electrode delivered stimulation at 70 μA, 40 Hz with varying PW (0%, 25%, 50%, 75%, 100% of MNA output; *n* = 10 cycles per condition) to evaluate gradual mechanoneural actuation.

### Biohybrid organ system

To further demonstrate the potential of MNA technology, we developed a biohybrid intestine system capable of modulating organ mechanics. This system consisted of a small intestine wrapped by the MNA, enabling control of intestinal contractions (squeeze and release) and offering potential applications in modulating organ-governed human physiology (e.g., appetite). Procedures for biohybrid intestine implementation in a rodent model are illustrated in Extended Data Fig. 5. SN access and MNA separation from MG followed prior methods. To simulate a filled intestine, diluted brown-colored saline was injected into the intestine. The distal end was tied securely, and a loop was formed by loosely tying the proximal end (∼2.5 cm from the distal knot). After filling, the proximal knot was secured, and the filled intestine was positioned near the posterior knee. The intestine was wrapped with the MNA, and the distal and proximal tendons were sutured together to form a circular configuration. Distributed surface marks were added to the intestine for video tracking. To evaluate gradual control of organ mechanics, a nerve cuff delivered varying levels of stimulation (70 μA, 40 Hz, 1–4% PW). Optical flow analysis (Matlab, MathWorks, USA) was performed on the recorded video to compute correlations between MNA actuation and organ movements. The velocity of the MNA and the targeted organ was smoothed using a moving average with a window size of 5 frames before the analysis.

### Statistics

All results were reported as mean ± standard error of the mean (SEM). Normality of the data was verified by the Shapiro-Wilk test (significance level *α* = 0.05). For normally distributed data, two-sided paired and unpaired *t* tests were used for within and between group comparisons, respectively. For data that violated normality, Mann–Whitney U tests were used for between group comparisons. The effects of the three time points on MNA mass and force production were analyzed using one-way analysis of variance (ANOVA). For the closed-loop force control performance analyses, slopes (*m*) between the target and output force values, *R*^2^, and *P*-values were reported. In the analysis of PMI performance, Kendall’s tau coefficients (τ) were reported to assess the relationships between residual muscle strains, evoked afferents, and MNA actuation levels, along with their corresponding *P*-values. In the analysis of biohybrid organ performance, RMS velocity values for the MNA and actuated organ, along with the Pearson correlation coefficient (*r*) and its corresponding *P*-value between the MNA and actuated organ, were reported. We performed statistical analyses using MATLAB 2022b (Mathworks, USA).

## Data availability

All data are available in the manuscript or Supplementary Information.

## Code availability

All data were collected through software available from Tucker-Davis Technologies (TDT) and Aurora Scientific.

## Author contributions

H.S., G.H and H.M.H. conceived the study concept. H.S., G.H., and G.N.F. contributed to surgical implementation. H.S. and G.H. performed overall actuator characterization, closed-loop control testing, and biohybrid system implementation and evaluation. H.S., G.H., and S.H.Y. contributed to nerve block design and implementation. S.G. assisted with muscle fatigue data collection. H.S. analyzed neuromechanical data and G.H., C.H and S.S. analyzed histological data. H.S. and G.H. prepared figures and wrote the original manuscript and H.S., G.H., H.M.H. edited the manuscript. H.M.H provided management and funding for the project. All authors revised and approved the final manuscript.

## Acknowledgements

We thank E. Boyden for sharing microscopy resources and the Koch Institute’s Robert A. Swanson (1969) Biotechnology Center for technical support, specifically the histology core for help in processing and imaging histology samples. Funding by the K. Lisa Yang Center for Bionics at MIT and the MIT Media Lab Consortia. G.H. is supported by a K. Lisa Yang ICoN fellowship and a CONACYT doctoral fellowship. G.N.F. is supported by the NIH/NINDS R25 program and the Neurosurgery Research Education Foundation and American Academy of Neurological Surgeons.

---

**Extended Data Fig. 1.**
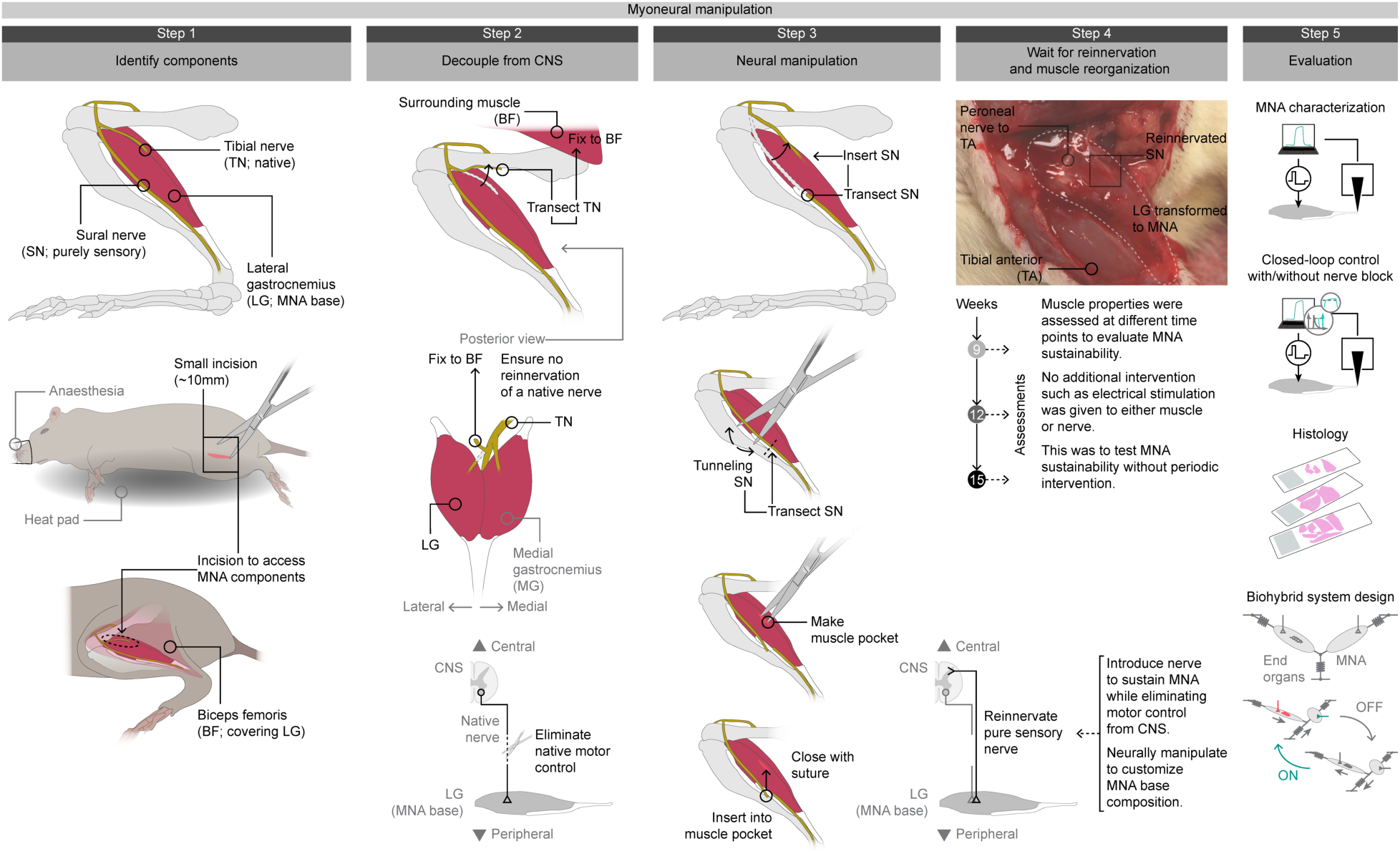
MNA design, implementation, and evaluation. Step 1: MNA design involves using the lateral gastrocnemius (LG) as the basis, with the sural nerve (SN; purely sensory) serving as a myoneural manipulator to enhance fatigue resistance. Each component is assessed through a small incision on the lateral side of the limb. Step 2: A procedure entails eliminating native motor control of the MNA basis by transecting a native motor nerve, specifically a branch of the tibial nerve (TN) connected to the LG. To prevent unintended reinnervation back to the LG, the transected TN branch is sutured onto the biceps femoris (BF), a surrounding muscle. Step 3: The distal end of the SN is transected and embedded into a muscle pocket created on the LG. Step 4: We allow time for the SN to fully reinnervate the MNA basis and promote myoneural reorganization. Step 5: We perform a series of tests to characterize the fatigue resistance and closed-loop control performance of the MNA with and without nerve block. Additionally, histology analyses are performed to verify the myoneural mechanism underlying fatigue resistant MNA. Multiple biohybrid systems are designed to demonstrate the potential impacts of MNA technology.

**Extended Data Fig. 2.**
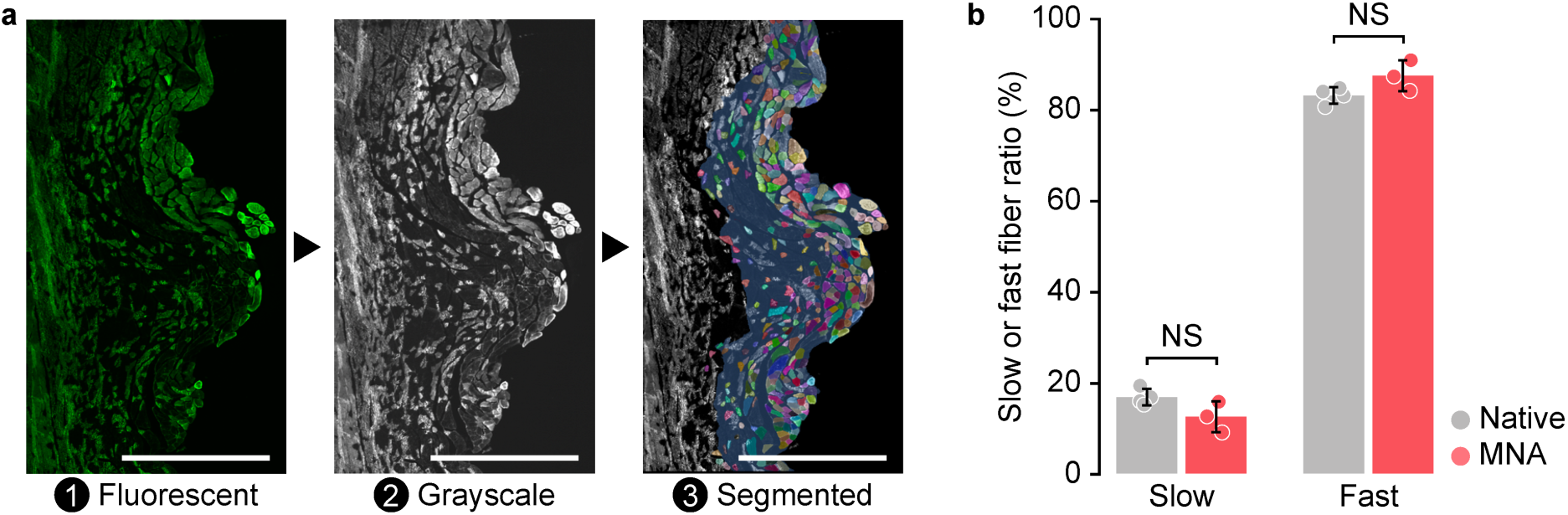
Histological analysis of MNA. **a**, Muscle tissue cross-sectional slides were stained using anti-myosin antibodies against slow and fast muscle fibers and images on a confocal microscope (1). Images were converted to grayscale before segmentation (2). Images were automatically segmented using Cellpose Cyto3 model. Pixels were then used to compute the total muscle and fluorescent area (3). Scale bar is 1 mm. **b,** Slow or fast fiber ratio percentage for native and MNA.

**Extended Data Fig. 3.**
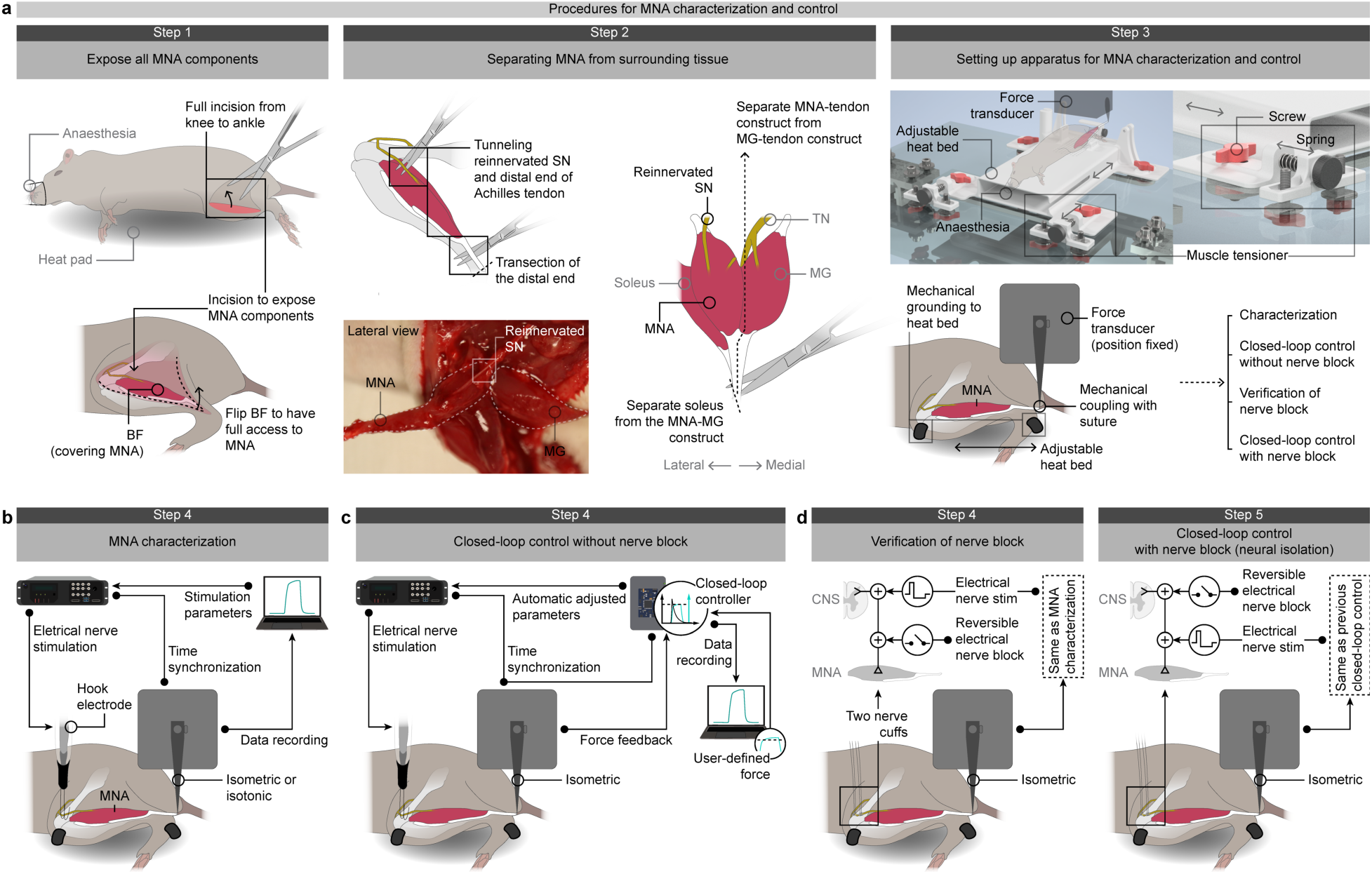
Experimental setup for MNA characterization and closed-loop control with and without nerve block. **a**, Common procedures for MNA characterization and closed-loop control are outlined. Step 1: A procedure involves making an incision extending from the knee to the ankle to fully access the MNA construct. Step 2: We clear out connective tissues surrounding the sural nerve (SN) to create space for placing a hook electrode or nerve cuffs. Subsequently, we proceed to separate the MNA construct from the medial gastrocnemius (MG) and soleus. Step 3: We position the prepared rat on an experimental apparatus comprising a heated bed with a muscle tensioner and a force transducer. We mechanically ground the patellar tendon and anterior talofibular ligament to the heated bed and connect the distal end of the MNA construct to a force transducer, while adjusting the position of the heated bed to pre-tension the MNA. **b**, Step 4: For MNA characterization, we apply a series of electrical nerve stimulations using a hook electrode placed on the SN while simultaneously recording MNA force outputs from the force transducer. **c**, Step 4: In the MNA closed-loop control without nerve block, force readings from the force transducer are fed to a controller to automatically adjust a stimulation parameter and to reach a target force value. **d**, Step 4: For the MNA closed-loop control with nerve block, two nerve cuffs are placed for MNA control and nerve block, respectively. We initially verify the nerve block by applying it distal to the electrical nerve stimulation of the MNA. By shunting neural signaling distal to electrical nerve stimulation, the MNA becomes non-responsive to nerve stimulation. Step 5: After verifying the nerve block, we swap the positions of the nerve block and stimulation, applying the nerve block proximal to the nerve stimulation. This shunts neural signaling proximal to nerve stimulation, providing neural isolation to the MNA from the CNS during actuation.

**Extended Data Fig. 4.**
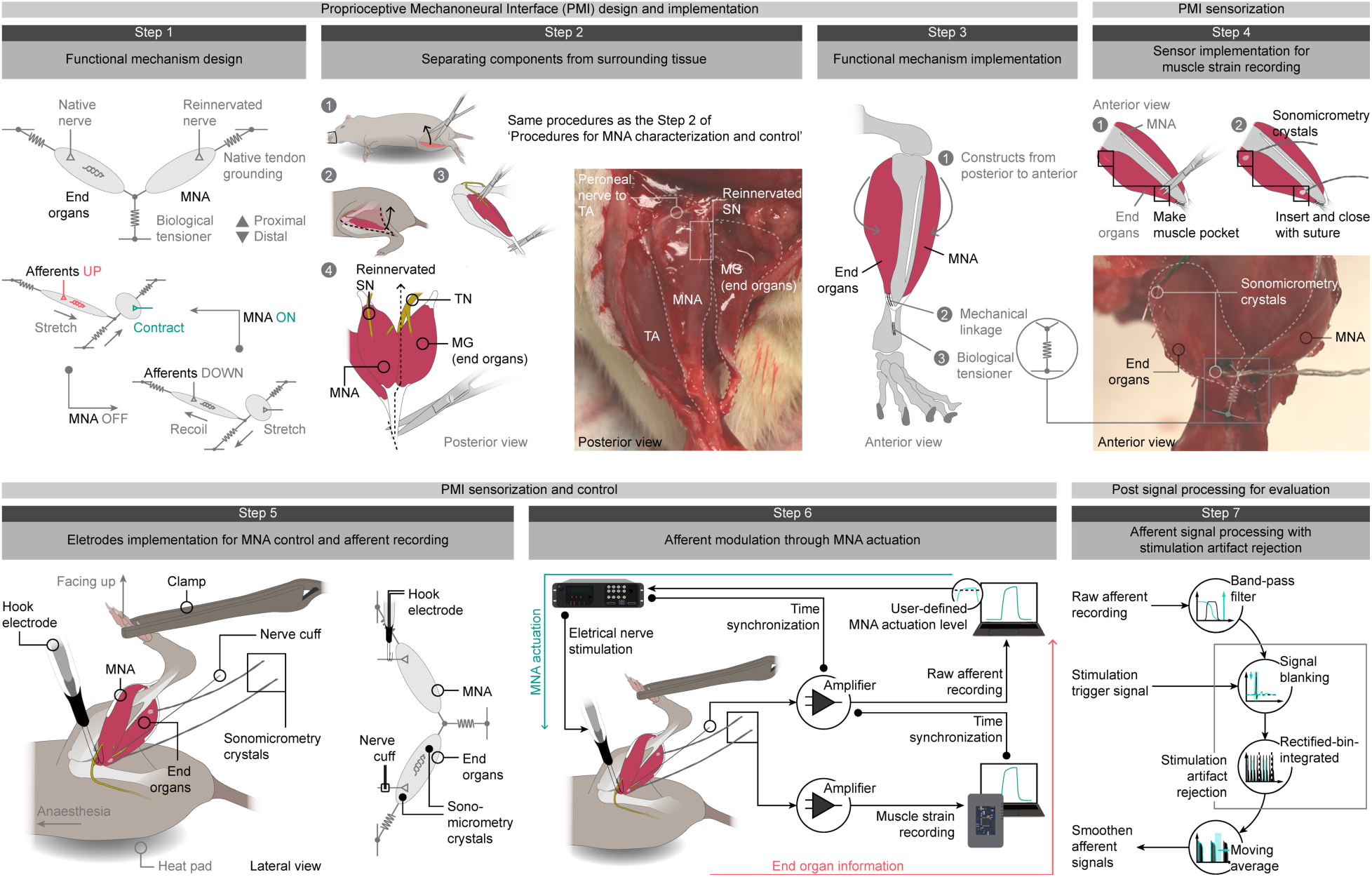
Proprioceptive Mechanoneural Interface (PMI) implementation and evaluation. Step 1: PMI functional mechanism consists of a MNA serially linked to a residual muscle (end organs) and a biological tensioner. Step 2: We separate the MNA construct from the surrounding tissue using procedures identical to those employed for MNA characterization and closed-loop control (see Extended Data Fig. 2a for more details). Step 3: We use the medial gastrocnemius (MG) as the target end organs, bringing both the MNA and end organs constructs from posterior to anterior for mechanical linkage, and anchoring the distal portion of the end organs tendon to the tibia. Step 4: We install a pair of sonomicrometry crystals into the end organs to track strain. Step 5: Nerve stimulation is delivered to the MNA through a hook electrode. Step 6: We perform varying levels of MNA actuation to assess the gradual afferent controllability. Step 7: We filter out stimulation artifacts and smooth the recorded afferent signals for further analysis.

**Extended Data Fig. 5.**
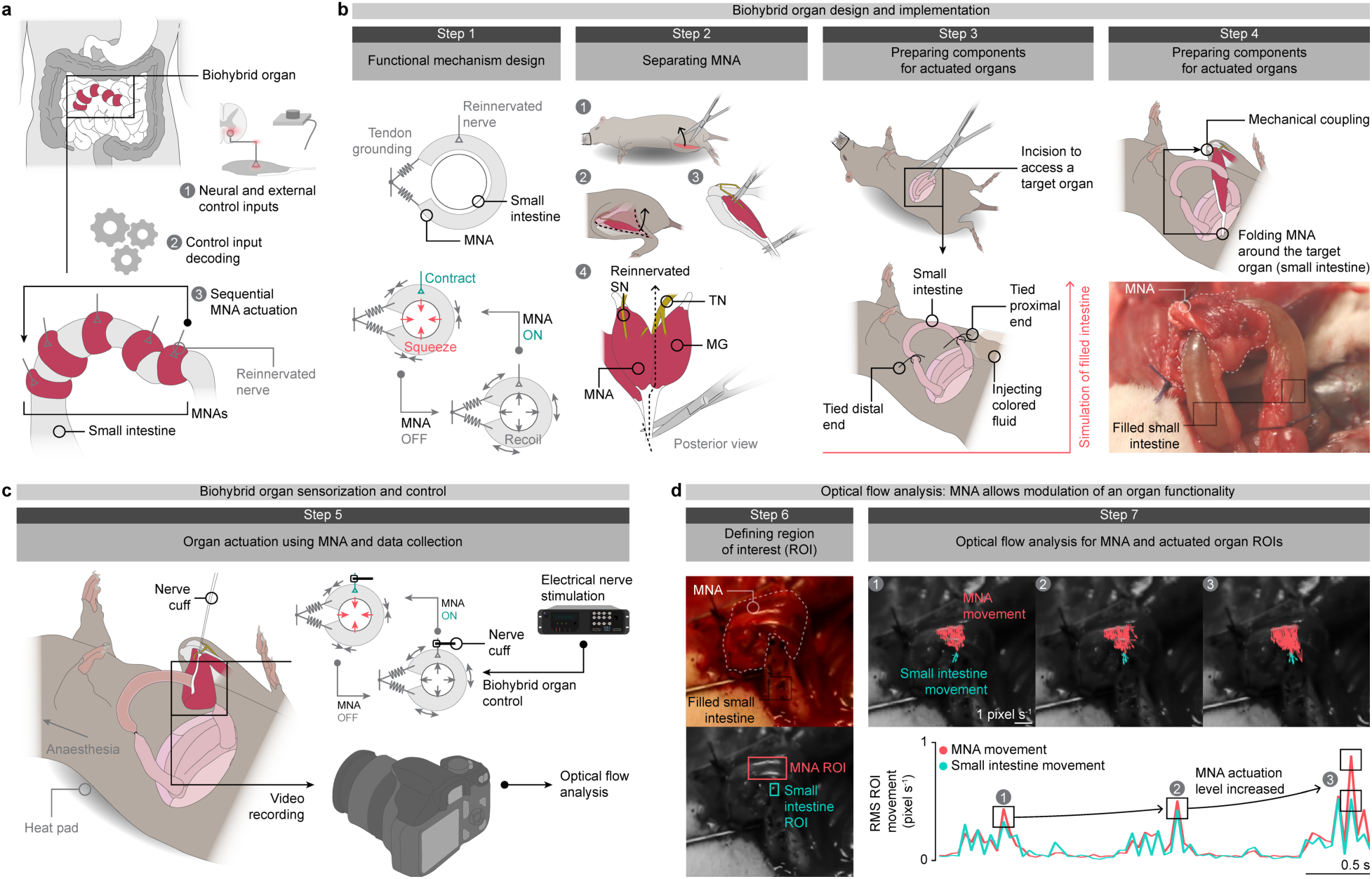
Organ mechanical modulation using a MNA. **a**, A conceptual diagram illustrates a biohybrid organ system designed for the small intestine. Neural signals originating from the CNS and peripheral nervous systems, as well as external signals, can serve as control inputs for the biohybrid organ system. **b**, Step 1: The biohybrid organ system comprises a MNA wrapped around the small intestine, aiming to emulate the mechanical functionality of the organ. Step 2: We separate the MNA construct from the surrounding tissue using procedures identical to those employed for MNA characterization and closed-loop control (see Extended Data Fig. 2a for more details). Step 3: We make an incision on the abdomen to access the small intestine. After tying the distal portion of the small intestine, we inject fluid to simulate a filled state and subsequently tie the proximal portion of the small intestine. Step 4: We then wrap the small intestine with the MNA. **c**, Step 5: Nerve stimulation for biohybrid organ modulation is delivered to the MNA through a nerve cuff. We record videos of the biohybrid organ modulation for further analysis. **d**, Step 6: The region of interest (ROI) for both the MNA and the actuated organ is selected for optical flow analysis. Step 7: The root-mean-square (RMS) velocity values for the MNA and actuated organ are plotted.

**Extended Data Fig. 6.**
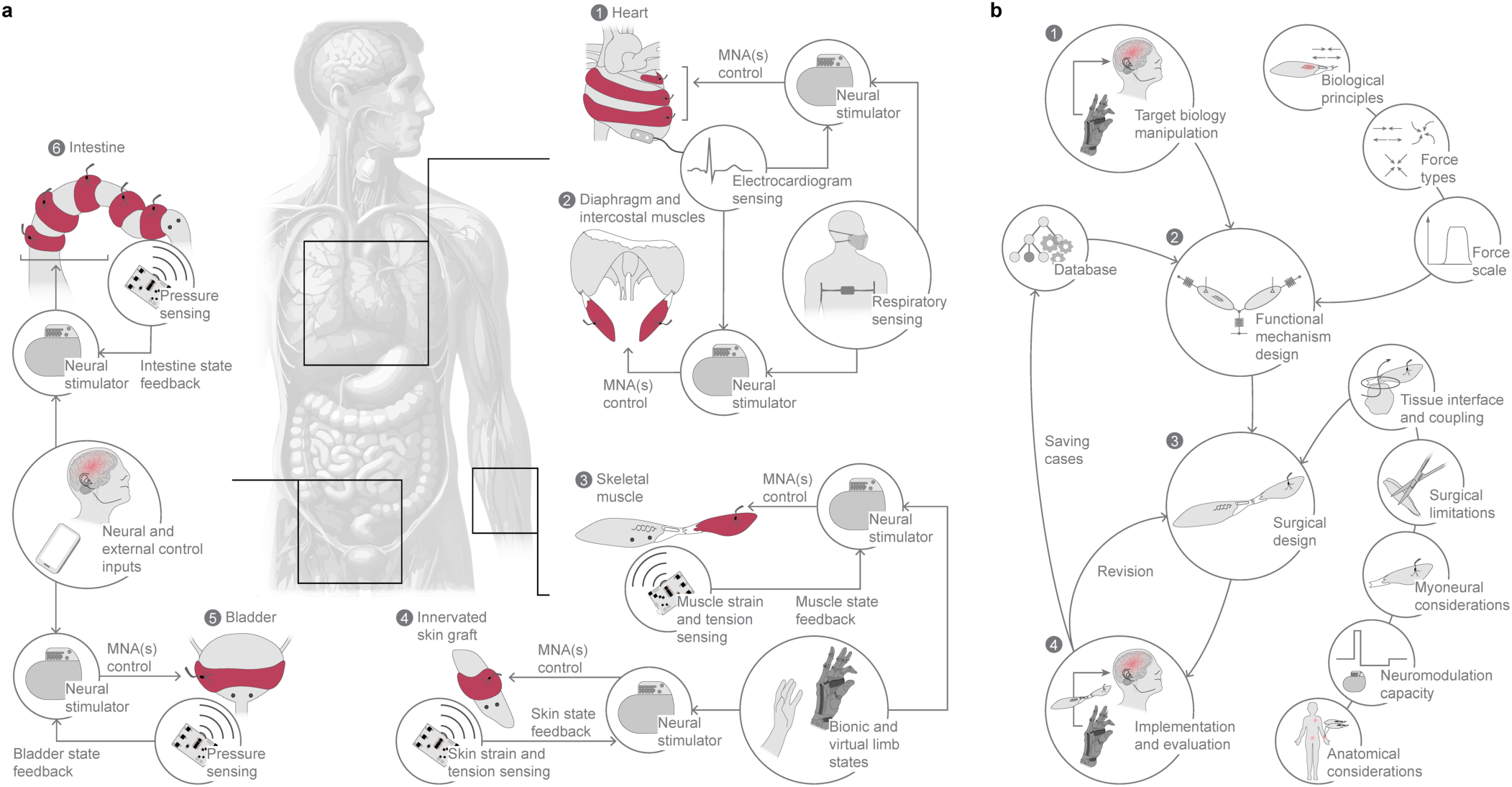
Design of biohybrid organ systems. **a**, The MNA enables actuation with full bi-directional isolation from the central nervous system (CNS), unlocking the creation of multiple biohybrid organ systems, from autonomic modulation to neural feedback. MNAs can be wrapped around the heart to provide mechanical actuation. Ventricular systole can be computed from electrocardiogram (ECG) leads and respiratory signals to synchronize nerve stimulation for closed-loop control of a MNA-based biohybrid heart (1). MNAs can be serially coupled to each of the diaphragm muscle attachment points and intercostal muscles to facilitate contraction during inspiration, which can be synchronized with respiratory and ECG signals (2). MNAs can be serially coupled to end organ muscles to modulate muscle dynamics for proprioceptive afferent signaling. Strain and tension sensing from end organ muscles through implanted sensors such as magnetic beads can be used to estimate muscle state feedback, while states from bionic and virtual limbs can be used to compute reference trajectories, and together, inform stimulation patterns for MNA closed-loop control (3). Similarly, MNAs can be wrapped around innervated skin grafts for cutaneous afferent signaling. Strain and tension sensing from skin can be used to estimate skin state feedback, while bionic and virtual limb states can be used to compute reference trajectories, and together, inform stimulation patterns (4). One or more MNAs can be wrapped around the detrusor muscle of the bladder to provide mechanical compression and assist in urinary control. Inputs from implanted sensors such as magnetic beads to measure bladder pressure, brain-computer interfaces and external control inputs such as smartphone commands can be used to inform stimulation patterns for closed-loop control of a MNA-based biohybrid bladder (5). A series of MNAs can be wrapped around the intestine to emulate peristaltic motion and modulate the gut-brain axis. Implanted sensors such as magnetic beads can estimate the intestine state coupled with neural and external control inputs for closed-loop MNA control (6). **b,** Biohybrid system design workflow using MNAs. The workflow begins with the target organ to be controlled (1). Considerations such as biological principles inform motion dynamics such as contractile and fatigue characteristics, attachment points, and tension levels. Force types guide required dynamics such as torsional, squeezing, peristaltic, or contraction. Force scale informs the magnitude and duration that is required to meet functional demands. Together, these elements define the functional mechanism design (2). After the mechanism design is completed, surgical design is performed based on considerations such as the tissue interface and coupling, for example, if the MNA needs to interact with cardiac tissue, a material that electrically isolates tissue would be required. Surgical limitations guide which tissue can be used to create the MNA. Myoneural considerations guide the type and size of muscle, sensory nerve, and number of each that would be required to meet functional demands. Neuromodulation capacity determines stimulation parameters to achieve desired functionality. Finally, anatomical considerations guide optimal locations to obtain MNA components and surgical approach to construct the biohybrid system (3). The surgical design is then implemented and evaluated. If required, a revision surgery can be performed to further optimize the system. Functional biohybrid systems are saved in a database that can later inform new mechanism designs (4).

